# Molecular Dynamic simulations of Aβ42 dimers with solid-state NMR restraints capture the key structural motifs in Aβ42 fibrillation pathways

**DOI:** 10.64898/2026.04.17.719214

**Authors:** Angelo L. Chu, Betty S. L. Chu, Wei Qiang

## Abstract

Formation of the β-amyloid (Aβ) plaques is a pathological hallmark of Alzheimer’s disease (AD), and is believed to be a primary cause of dementia in elderly individuals. In the present work, we simulated the conformational evolution of Aβ42 dimers in solution and in membrane-like environment to explore the folding of Aβ42 along fibrillation. The molecular dynamics (MD) simulation was steered by experimental internuclear distance restraints obtained using solid-state nuclear magnetic resonance (ssNMR) spectroscopy. Our results revealed that several hydrophobic and polar motifs within the Aβ42 sequence played key roles in the early-stage nucleation process of fibrillation and those motifs are also the stabilizing agents in the mature fibrils judged by the energy contribution. Our results also indicated that the peptide association with membrane bilayers could modulate the structural evolution pathways towards fibrillation. These findings contributed to a better understanding of the molecular level structural polymorphisms inherent to Aβ42 fibrils. Further, the current work demonstrated that the combination of MD simulations with ssNMR-based experimental restraints provided a reliable method for studying structural changes of Aβ.

**Highlight:**

- Using solid-state NMR restraints guided molecular dynamic simulation, β-amyloid dimers displayed consistent β-strand-prone regions, which are major stabilizing segments for mature fibrils.
- β-amyloid dimers evolved differently with or without interacting with the lipid bilayers.
- Experimental restraints guided simulation provided molecular level insights about early-stage interactions along the progress of β-amyloid fibrillation

## 1. Introduction

Alzheimer’s disease (AD) is the most common cause of dementia. According to the Alzheimer’s Association (https://www.alz.org/alzheimers-dementia/facts-figures), seven million Americans are living with AD in 2025. The current cost of treatment of AD and other dementias amount to $384 billion and will approach $1 trillion by 2050. Pathologically, AD is marked by abnormal plaques and tangled fibers deposited in the patients’ brain^1, 2^. The plaques are composed of amyloid fibrils formed by a short peptide with 37-44 residues, called β-amyloid peptide (Aβ). Researchers have revealed the molecular structures of Aβ fibrils, containing multiple Aβ subunits bonded by interstrand N-H^…^O=C hydrogen bonds forming β-sheets^3^. While the elucidation of stable Aβ fibril structures partially explains the challenges in the clearance of senile plaques in AD patients’ brains, many questions remain unanswered. For example, Aβ fibrils possess significant molecular-level structural diversity, known as the structural polymorphism^4, 5^. Previous solid-state nuclear magnetic resonance (ssNMR) spectroscopy and cryogenic transmission electron microscopy (cryoTEM) studies unraveled the presence of such polymorphisms within the fibrils extracted from postmortem AD patients’ brain tissues^6, 7^. However, correlations between the polymorphic fibrils and these patients’ symptoms and clinical histories have not been established. Furthermore, extraordinary efforts have been made to develop therapeutic strategies for AD, among which the designing of antibodies that specifically target early-stage Aβ oligomers and protofilaments have proven promising. For example, Lecanemab, an FDA-approved immunotherapy drug, clears the plaques and slows the progression of early-stage AD^8, 9^. The rationale is that the Aβ oligomers and protofilaments represent the most toxic species that would damage the brain nerve functions, and targeting those species for clearance and removal would delay AD development^10^. Understanding the molecular level conformational evolution within the early-stage Aβ oligomers and protofilaments will provide crucial insights to address these current questions in both fundamental structural studies and therapeutic developments. However, unlike mature fibrils that possess well-defined structures, the molecular structural characterizations of early-stage Aβ intermediates are experimentally challenging due to their transient and disordered nature. Tycko and co-workers have characterized the early events in Aβ self-assembly using time-resolved ssNMR with rapid-mixing and freeze-trapping apparatus and dynamic nuclear polarization (DNP)^11, 12^. Their results demonstrated the development of local β-strand conformations and intra-strand contacts within 1ms, and the appearance of inter-strand contacts within 500ms^13^.

On the other hand, molecular dynamic (MD) simulations have been commonly utilized to investigate the molecular processes in silico, suitable to study the early events of molecular assembling. Previous MD studies have been carried out to understand Aβ aggregation from monomeric states both with and without model membranes^14–16^. While some simulations employed structural motifs derived from mature fibrils as their starting conformation^15^, with a few utilizing populated Aβ monomer or dimer conformations from pre-equilibrium conditions^14^, all-atom simulation in combination with coarse-grain simulation can only extend the simulation in milliseconds range, barely entering the time regime of the freeze-trapping ssNMR spectroscopy^13^. Therefore, knowledge gaps remains despite advances in computational power and experimental techniques.

We have recently investigated the molecular structural features of Aβ intermediates in the presence of membranes, focusing on the primary nucleation phase of fibrillation processes^17,18^. Using ssNMR spectroscopy with selectively labeled 40- and 42-residue Aβ (Aβ40 and Aβ42 respectively), residue-specific inter-strand and peptide-lipid distances were determined at intermediate incubation time points ranging from 1 to 15 h. Compared with the less pathogenic Aβ40, Aβ42 exhibits a higher overall lipid-proximity population. DNP-ssNMR further confirmed local secondary structural convergence and the presence of long-range tertiary contacts between the two main hydrophobic segments during the structural evolution process. In the current simulation, we attempt to incorporate these experimental restraints into the generation of initial Aβ42 conformations and the establishment of force fields. Our MD simulation will provide direct visualization of how Aβ42 oligomer structures evolve along the nucleation process in the presence of membrane, with the guidance of experimental restraints.

## 2. Experimental Section

### 2.1 Setup of the Simulated Systems

Aβ42 sequence (DAEFRHDSGYEVHHQKLVFFAEDVGSNKGAIIGLMVGGVVIA) was obtained fromGenScript at: https://www.genscript.com/peptide/RP10017-_Amyloid_1_42_human.html. A python code was used to generate the random structures of Aβ42 peptide as a starting PDB structure. ChimeraX was used to add any missing H atoms to the PDB. All structures and MD simulation were visualized and examined using VMD (Visual Molecular Dynamics) 1.9.4^19^. The all-atom MD simulations were performed using NAMD 2.14 (University of Illinois at Urbana–Champaign, http://www.ks.uiuc.edu/Research)^20^. CHARMM-GUI (http://www.charmm-gui.org) was used to generate a membrane lipid bilayer system with POPC^21^. Charmm3.6m force field and TIP3P water were employed in all the MD simulations. H_2_PO_4_^-^ 3.8mM, HPO_4_^2−^ 6.2mM were added to the system and sodium ions were also added to balance the charge. All runs were carried out at 310 K, pH 7.4. The system was equilibrated under isobaric–isothermic conditions with a pressure of 1.01325 bar. Periodic boundary conditions were set in all directions. Both the van der Waals and Coulomb force cutoffs were set to 1.2 nm in real space. The particle mesh Ewald (PME) method was applied for calculating the electrostatic interactions. The simulation was run at NMRbox (https://nmrbox.nmrhub.org/), an online platform hosted by the University of Connecticut providing supercomputing capability^22^.

### 2.2 Simulations in water environment

Six Aβ42 molecules were first put in a water box with the size 20.5×19.7×13.6 nm^3^ (531373 atoms), then energy minimized and equilibrated. The molecule positions were manually adjusted to form 3 dimers. The distance restraints for the dimers were applied using harmonic walls with the force constant increased gradually from 0.5 to 15.0 kcal/mol/Å^2^ to avoid the system crash at the beginning. The distance restraints were obtained from our original experimental measurement using solid-state NMR^18^ (Table 1, restraints obtained after 3 hours upon the initiation of Aβ42 assembling). LowerWalls and upperWalls were used to set the lower distance and upper distance limits for each restraint, respectively. After additional energy minimization and equilibration, the production ran lasted for 74 ns. The force constant for the distance restraints was 15.0 kcal/mol/Å^2^ during the production run.

**Table 1.** The distance restraints used in the simulation. Atom 1 and Atom 2 are the corresponding atoms in two Aβ42 molecules forming the same dimer. Three dimers are all restrained. There are no distance restraints between the dimers.

| Atom 1<br>(molecule<br>1) | Atom 2<br>(molecule<br>2) | Distance<br>(Å) |
| --- | --- | --- |
| A2-Cβ | A2-Cβ | >9.0<sup>1,2</sup> |
| F4-C' | F4-C' | >9.0 |
| G9-Cα | G9-Cα | >9.0 |
| V12-C' | V12-C' | 7.8-9.4 |
| F19-C' | F19-C' | 5.4-6.2 |
| A21-Cβ | A21-Cβ | 5.8-6.6 |
| G25-Cα | G25-Cα | 5.8-6.6 |
| A30-Cβ | A30-Cβ | 5.7-6.3 |
| G33-Cα | G33-Cα | 6.8-8.0 |
| M35-C' | M35-C' | 5.8-6.6 |
| G38-Cα | G38-Cα | 6.3-7.7 |
| A42-Cβ | A42-Cβ | 6.3-7.3 |

<sup>1</sup>Distance restraints obtained from Ref. 15, through <sup>13</sup>C-PITHIRDS-CT spectroscopy.
<sup>2</sup>A restraint > 9.0 Å indicated farther interstrand distances beyond experimental detection limit.

### 2.3 Simulations in the presence of POPC membrane

Two Aβ42 dimer structures were taken from the above simulation in water and placed in the upper water layer of a membrane system. One Aβ42 dimer was set to be partially inserted into the POPC bilayers from V24 to L34. This configuration was selected based on our ssNMR distance restraints^18^ between specific residue and phospholipids headgroup (Table 2), where the A21-G33 segment is in close proximity with membrane ^31^P, whereas F19 or M35 exhibit longer distances (> 9Å). Although explicit distance values were not applied as restraints with enforced force constants, a reasonable starting configuration was manually adjusted to be consistent with the experimental observations. The second Aβ42 dimer was set to be slightly inserted the bilayer at its C-terminus (at Ile 41and A42 in one chain (protein C) and from G38 to A42 in the other chain (protein D)), as a comparison to the first configuration. The minimum water height on top and bottom of the system was set at 22.5 Å during CHARMM-GUI setting, resulting in a box size of about 13.2×13.2×13.2 nm^3^, containing 205305 atoms. The energy minimization and equilibrium were run in steps by first releasing the constraints on the lipid acyl chains while fixing the lipid headgroups and the protein, then releasing the constraints on the lipids while keeping the proteins fixed, followed by only fixing protein backbones. Finally, the constraints on the protein backbones were released and the distance restraints in table 1 were applied with a force constant of 25.0 kcal/mol/Å^2^. Other settings were the same for the equilibrium and the production, as listed in 2.1 section. The production ran lasted for 120 ns.

**Table 2.** The distance measured using ssNMR between residues in Aβ42 molecules and lipids headgroup ^31^P atom.

| Atom 1<br>(Aβ42) | Atom 2<br>(lipids) | Distance<sup>1,2</sup><br>(Å) |
| --- | --- | --- |
| A2-Cβ | $^{31}\text{P}$ | 6.6-7.0 |
| F4-C' | $^{31}\text{P}$ | >9.0 |
| G9-Cα | $^{31}\text{P}$ | 6.9-7.1 |
| V12-C' | $^{31}\text{P}$ | >9.0 |
| F19-C' | $^{31}\text{P}$ | >9.0 |
| A21-Cβ | $^{31}\text{P}$ | 7.3-7.9 |
| G25-Cα | $^{31}\text{P}$ | 6.6-7.1 |
| A30-Cβ | $^{31}\text{P}$ | 5.8-6.3 |
| G33-Cα | $^{31}\text{P}$ | 5.7-6.3 |
| M35-C' | $^{31}\text{P}$ | >9.0 |
| G38-Cα | $^{31}\text{P}$ | >9.0 |
| A42-Cβ | $^{31}\text{P}$ | 4.4-5.6 |

<sup>1</sup>Distance restraints obtained from Ref. 15, through $^{13}\text{C}$ - $^{31}\text{P}$ REDOR spectroscopy.
<sup>2</sup> A restraint > 9.0 Å indicated farther interstrand distances beyond experimental detection limit.

### 2.4 Analysis of Aβ42 dimers and the published Aβ42 fibril structures

Although Aβ42 was started with random coil conformations, the peptide developed several secondary structure elements during the structural evolution. The secondary structure of each Aβ42 residue in the dimers was computed using STRIDE (**Str**uctural **ide**ntification)^23^ incorporated in VMD. Total backbone RMSD for each Aβ42 vs time was also recorded for the total production run period in comparison to the first frame in the production run. For the structural evolution in the presence of lipids, salt bridge analysis was also carried out for the dimers using the function in VMD, recording any ion-pairs with oxygen-nitrogen distance within 3.2 Å. A tcl script was applied to access the intermolecular contacts within a dimer, which would record any atom pairs between proteins within 4.5 Å. All recorded distances for a residue pair were averaged in calculation of the intermolecular residue distance. Distance between F19 and I32/L34 within a dimer was accessed using center of mass. During the MD simulation in the water box or in the presence of POPC bilayers, different Aβ42 dimers did not come to close contact with each other. Therefore, no intermolecular contact between dimers was accessed. Since not many interactions were observed between the lipids and one of the Aβ42 dimers, and the other Aβ42 dimer inserted in the membrane also did not perturb the membrane much, no analysis on the POPC lipid bilayers were done.

FoldX^24^ was utilized to calculate the energy contribution of each residue to the total free energy of the fibril (https://foldxsuite.crg.eu/products#foldx). All Aβ42 fibril structures were downloaded from pdb databank (https://www.rcsb.org/) and were modified to contain 5 layers of cross-β unit prior to analysis. Structures were first subjected to side-chain energy minimization using the FoldX RepairPDB command. This procedure optimizes side-chain conformations and selects the lowest-energy rotamer for each residue from a rotamer library. Then the FoldX SequenceDetail function was applied to compute per-residue free energy contributions (ΔG) and the associated thermodynamic components. To avoid boundary artifacts, subunits located at both ends of the extended fibril were excluded from analysis. Therefore, only the middle three layers were used to generate the data. The heatmap was generated using a python script.

## 3. Results and Discussion

### 3.1 Construction of the initial Aβ42 dimers with experimental restraints and simulations in solution

Three Aβ42 dimers were constructed from six Aβ42 molecules with restraints enforced within each dimer during the MD simulation. No restraints between the dimers were applied to allow the dimer to move freely in the system. Aβ42 sequence possesses mostly charged/polar residues and mostly hydrophobic/nonpolar residues at its N- and C-terminal halves, respectively. The ssNMR-based internuclear distances restraints (Table 1) were adopted from the Aβ42-membrane intermediates with three-hour incubation, and well spread from the N-terminal A2 to the C-terminal A42. For the first three N-terminal residues (A2_Cβ, F4_C’ and G9_Cα), no experimentally derived inter-strand distance was available, and a 9 Å low-boundary distance restraint was applied, as it represented the upper limit of ssNMR measurements.

Analyses of the secondary structures of dimers during the structural evolution in the water box showed that certain segments, such as K16-F20, had high tendencies to form β-strand conformation (Figure 1 and Supplement Figure S1) even without the backbone conformational restraints (e.g., dihedral angles). Two out of the three dimers showed this stabilized β-strand-like motifs while the third one remained mostly random coil. Furthermore, dimer 1 developed a U-shaped domain with F19-20 and I31-32 facing each other, and dimer 2 adopted a stretch of β-strand at ^30^AII^32^. These secondary and tertiary structural features were commonly observed in mature Aβ fibrils, suggesting that their formation at earlier stages such as the nucleation phase may be important to produce effective elongation-prone nuclei. It was noted that dimer 3 did not exhibit a stable β-strand motif during the production run and may require a longer simulation time to form such a structural motif. The diversities in structural evolutions may be due to the different starting conformers, all of which were set to be random at the secondary structure level. The two dimers with stabilized β-strand motifs (Figure 1) were used as the starting structures for the next round of MD simulation in POPC membranes.

**Figure 1.**
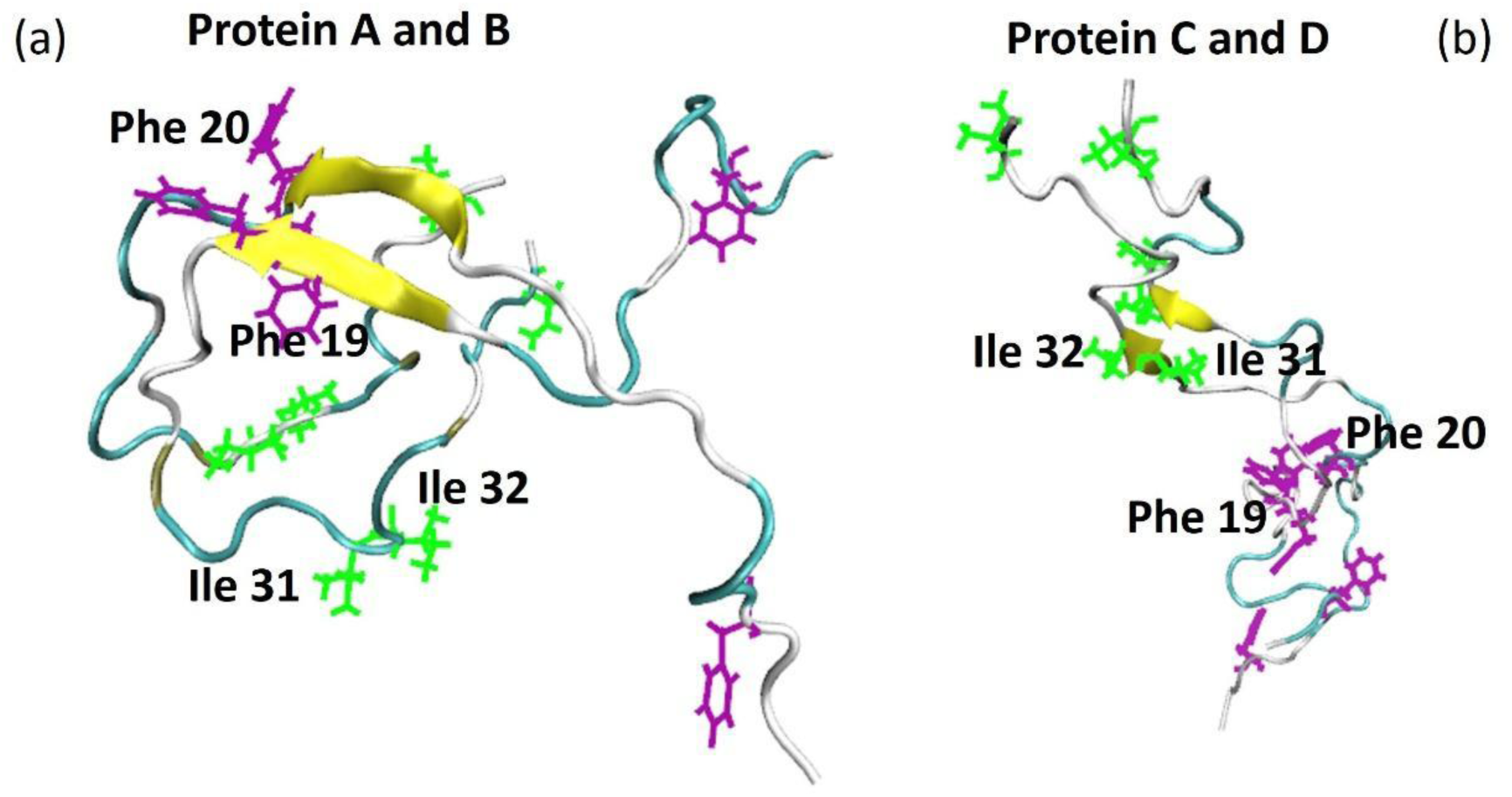
The selected dimer 1 (a, protein A and B) and dimer 2 (b, protein C and D) structures that were used in the next step, the simulation carried out in the presence of POPC lipids. Purple residues are Phe while green residues are Ile. Dimer 1 displayed β-strand structure around Phe19-20 and dimer 2 displayed β-strand structure around Ile31-32.

### 3.2 Simulations in POPC membrane

Using CHARMM-GUI, a membrane system was constructed with dimer 1 inserted in the lipids from V24 to L34 and dimer 2 slightly inserted in the lipids from G38 to A42 in one chain (protein D) and at Ile 41and A42 in the other chain (protein C) (Figure 2a). Dimer 1 with a U-shaped conformation was positioned relatively deep within the membrane as this insertion depth and orientation were consistent with our experimental restraints^18^. Dimer 2 with an extended conformation cannot be manually inserted into the bilayer without violating the experimental restraints; therefore, it was randomly positioned on the membrane surface as a comparison. The ssNMR-based inter-strand distance restraints (Table 1) within each dimer were maintained throughout the simulation using kinetic force, whereas the position of the dimer relative to the membrane was not restrained, allowing assessment of how the membrane influences dimer configuration. During the 120 ns production run, dimer 1 was never released from the membrane, although at the end a major part of the buried residues had come out of the membrane. Dimer 2 quickly moved out of the membrane at the beginning of the production run and maintained in the water phase (Figure 2b, c).

**Figure 2.**
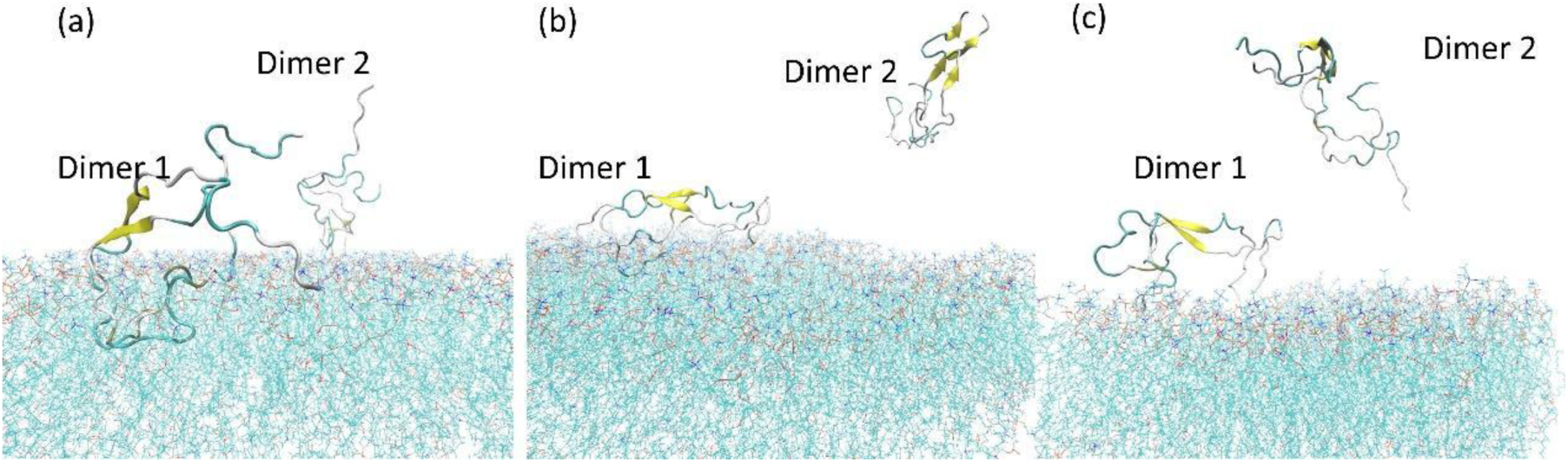
The structural snapshots of 2 dimers in the membrane system. (a) at the beginning; (b) at the 70ns time frame; (c) at the 120ns time frame.

Since Aβ42 contained many charged residues at its N-terminal sequence, salt bridges were formed frequently during the MD run, rendering turns and loops forming transiently (Figure 3). Interestingly, dimer 1 had significantly fewer salt bridges formed compared to dimer 2. Many salt bridges in dimer 2 also crossed longer sequences, such as between E3 and K28, R5 and E22 or D23, D1 and K16. The observation can be explained by the fact that dimer 1 was restricted by the membrane bilayers while dimer 2 had more freedom to explore the conformational space. This was also supported by the secondary structure analysis for dimer 1 (Supplementary Figure S2) and dimer 2 (Supplementary Figure S3). The results showed that dimer 1 maintained the β-strand at ^17^LVFF^20^ (protein A and B in Figure 4), but no additional β-strand motif developed. On the contrary, dimer 2 developed additional short stretches of β-strand with the time evolved (shown in blue in Supplementary Figure S3). Besides the initial β-strand motif around I31-32, it gained β-strand structures at F17-20, S25-N27 and M35-V40, representing hotspots for adopting β-strand conformation (protein C and D in Figure 4).

**Figure 3.**
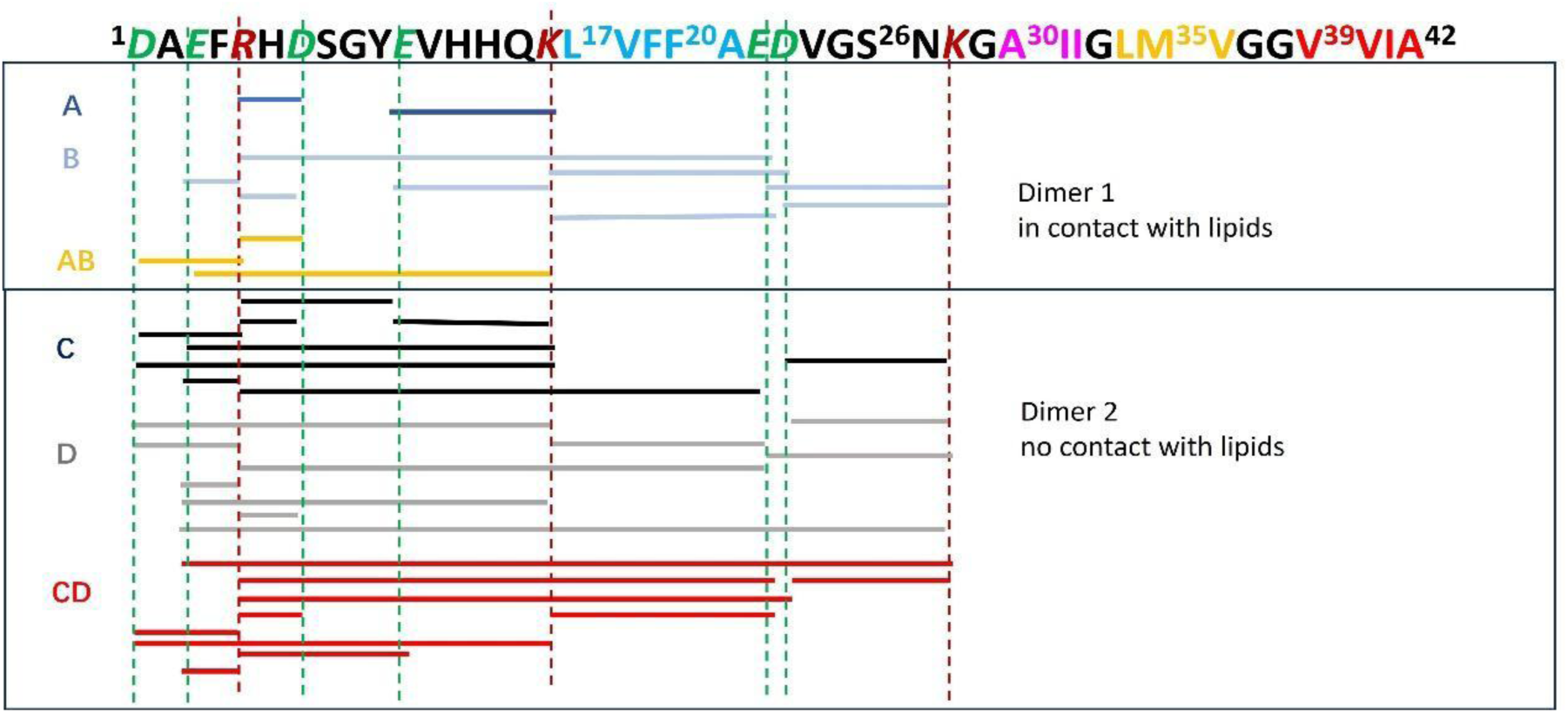
Salt bridges analysis using VMD, recording any ion-pairs with oxygen-nitrogen distance within 3.2 Å. The Aβ42 sequence was also displayed above the figure, showing negative charged residues in green and positive charged residues in marron. Small stretches of hydrophobic residues were also highlighted using colors. Salt bridges were indicated by short lines connecting green and marron residue pairs. In dimer 1, salt bridges in protein A were shown in dark blue, while salt bridges in protein B were shown in light blue. Salt bridges between a protein A residue and a protein B residue were shown in yellow. In dimer 2, salt bridges in protein C were shown in black, while salt bridges in protein D were shown in grey. Salt bridges between a protein C residue and a protein D residue were shown in red.

**Figure 4.**
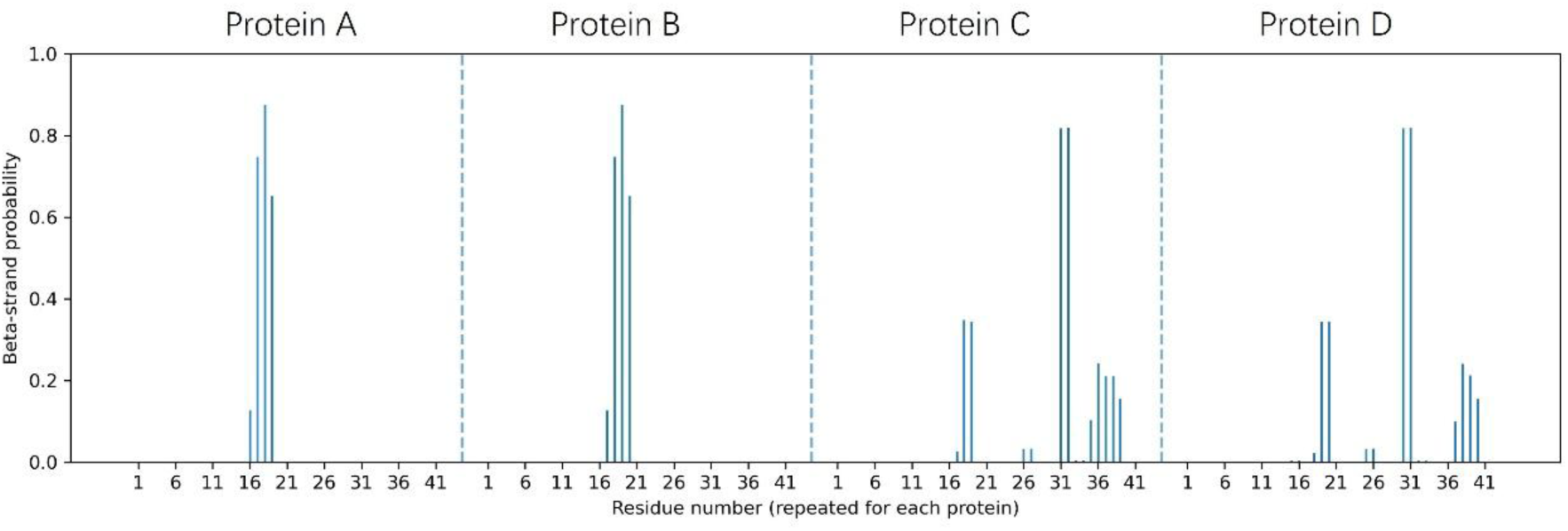
The probability of β-strand formation during the structure evolution. The probability was calculated based on data in Supplement Figure S4, and S5. Only “E” was considered as β-strand conformation. Dimer 1 formed by protein A and B maintained β-strands at around residue 16-20 in POPC bilayers. Dimer 2 formed by protein C and D had 4 stretches of β-strand at residue 17-20, 25-27, 31-32, 35-40.

It is worth noting that the observed β-strand hotspots (underlined, top sequence, Figure 5) through MD simulation were close to residues with ssNMR-based experimental inter-strand distance restrains (highlighted in yellow, bottom sequence, Figure 5), but not exactly overlapped. For example, V12, A21, A30, G33 and A42 were experimentally restrained, but were not shown as the β-strand hotspots in simulation. In addition, the ssNMR-based restraints on these residues were set with low boundaries longer than 5.4 Å, beyond the 4.8 Å inter-strand distances in typical extended β-sheets. Therefore, our observation of β-strand formation within these hotspots were likely to be driven by intrinsic properties of individual residues and the Aβ sequence, not an artificial effect due to the application of experimental restraints.

**Figure 5.**
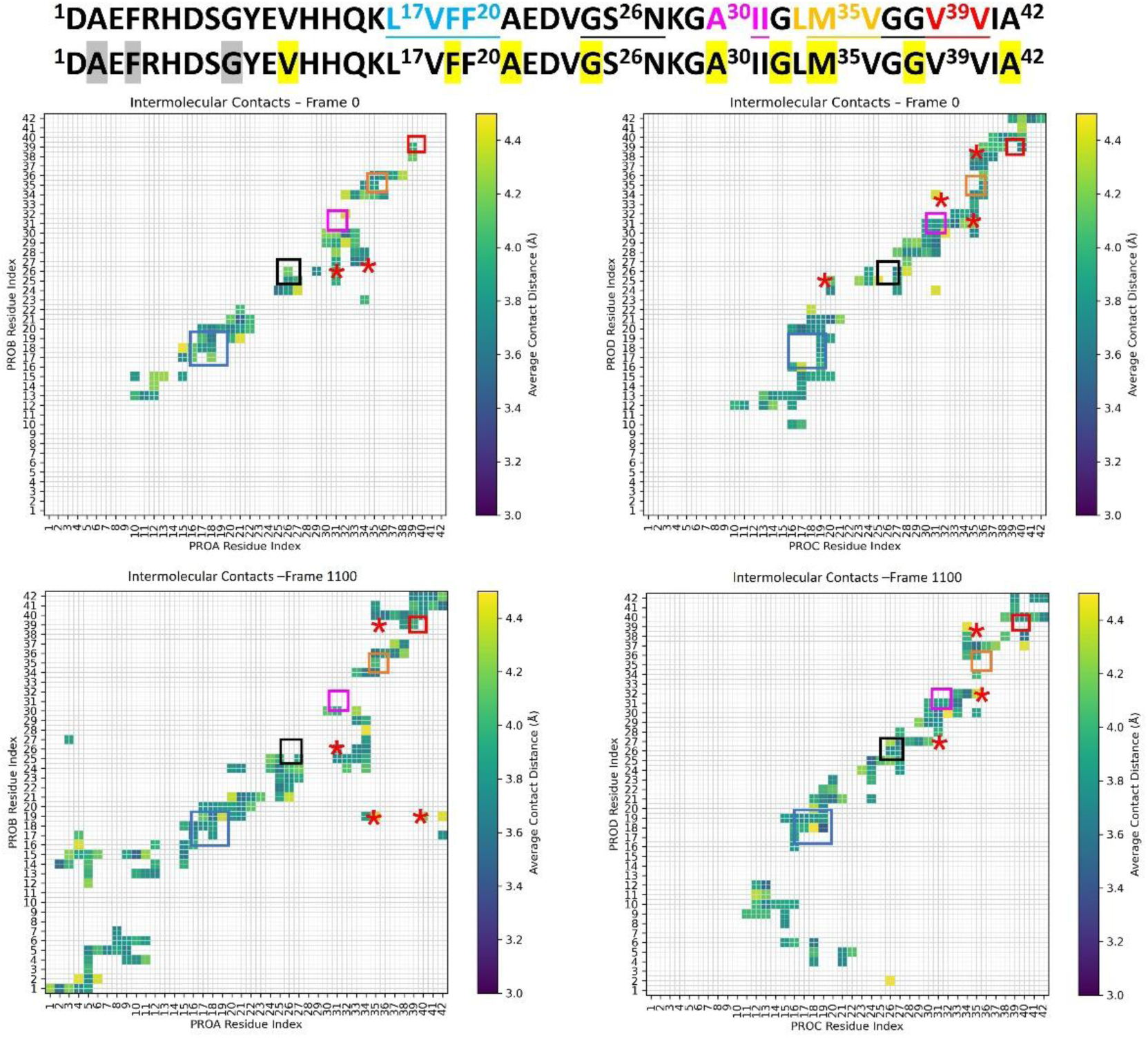
The intermolecular contacts within a dimer. Proteins A and B are two chains for dimer 1, and Proteins C and D are two chains for dimer 2. The four hotspots for β-strand were underlined in the sequence, also marked using squares in the contact plots. The last hotspot residue 35-40 was split into two segments: residue 35-36 and residue 39-40. The red star indicated the existence of contacts between residues in marked squares.

To explore the time-dependent tertiary structural changes in the two dimers, we plotted the intermolecular contacts within 4.5 Å at simulation time zero and at 110 ns (Figure 5). This displayed distance in figure 5 is the averaged value accounting to all < 4.5 Å distances of the corresponding residue pairs. The four hotspots of β-strand motifs from the secondary structure analysis were underlined in the sequence, and marked with squares in the contact plots. At time zero, no interstrand contacts were found between residue 1 to 10 for both dimers, consistent with the ssNMR-based restraints where residues A2, F4 and G9 showed > 9 Å interstrand distances. In addition, several blank spots along the diagonal line (e.g., E22-G25 in both dimers) were observed, suggesting the presence of highly disordered initial structures for both dimers with non-uniform interstrand proximity. The comparison between the contact map at 110 ns with time zero showed that there were more inter-strand proximities between the highlighted segments in dimer 1, marked in red stars. For instance, the segment around F19 (blue square, Figure 5) showed close contact with the segments around L34 (yellow square) and V40 (red square). The approaching between F19 and I32/L34 in dimer 1 was further supported by the analysis of these specific inter-residue distances through the simulation in POPC (Supplement Figure S4). Interestingly, these features were absent in dimer 2 at 110 ns, which still adopted extended conformation within its C-terminal sequence and did not exhibit pronounced difference compared to its initial contact map. There were contacts only between adjacent β-strand-prone segments in dimer 2. In addition, close diagonal contacts were observed in dimer 1 for the first 10 N-terminal residues.

Two conclusions can be drawn from the comparison between dimer 1 and dimer 2 in POPC bilayer, in the context of early-stage conformational changes: First, the contact between Aβ dimer and membrane restrict the flexibility of possible secondary structures that can be adopted by the peptide backbone. It was seen that dimer 2 (in solution) possessed much more transient secondary structures at multiple short motifs compared to dimer 1 (associated with the bilayer). Second, at the tertiary structure level, the involvement of membrane significantly prompts the formation of long-range contacts within the dimer and the extension of inter-strand proximity towards the disordered N-terminal segment. Combining these two insights, our simulation indicates that early-stage Aβ dimers may take distinct structural evolution pathways in the presence or absence of membranes, and the involvement of membranes benefits the production of elongation-prone nuclei.

Aβ oligomers are important targets for developing diagnostic and treatment agents for AD, and are also key intermediates for understanding the amyloidogenic misfolding process of Aβ. However, deciphering the structural properties for early-stage Aβ oligomers are challenging. Application of traditional MD approach to this problem would require simulations beyond multi-microsecond timescale to observe structural folding and the development of β-strand conformational motifs^14^. Alternatively, experimentally derived conformations from mature fibrils could be employed as initial conformers to accelerate the simulation. However, these structures may not directly represent the features in early-stage intermediates under experimental conditions^15^. Current MD simulation employed ssNMR-based inter-strand distance restraints obtained on early-stage Aβ_42_ intermediates, for constructing and modeling dimers. We showed that these dimers developed β-strand secondary structure as well as experimentally relevant tertiary contacts within ~ 100 ns both in the absence or presence of lipid bilayers. Therefore, our study demonstrated the feasibility of performing ssNMR-experimental-restraint-based MD simulations with higher computing efficiency.

### 3.3 β-strand-prone motif

While the charged residues located mainly at the N-terminal half of the sequence formed transient salt bridges (Figure 3), we observed that β-strand motifs with the dimers mostly occurred around the hydrophobic residues and polar residues towards the C-terminal segments (Figure 4). We showed that L17-F20 was a highly conserved β-strand-prone motif in both dimers and in either solution or membrane-like environments, consistent with previous studies showing that F19-20 initiated the β-strand folding in amyloid fibrils through side-chain π-π interactions^25–27^. More short motifs towards the C-terminus, including G25-Q27, I31-I32, M35-V36, and V39-40, were identified as β-strand-prone. The hydrophobic segments are commonly involved in the β-sheet cores in mature fibrils^28^. The S26-Q27 motif was also frequently found in other amyloid sequences^29^. Therefore, we hypothesize that these hydrophobic and polar motifs are hotspots for the β-strand formation, and the consequent short β-strands further fold into various tertiary structures depending on the molecular environments of fibrillation. The variations in the tertiary association of these short motifs may underline the well-known structural polymorphisms in mature Aβ fibrils^4^. Figure 6 displayed the stabilizing energy of each residue in Aβ42 sequence for a list of published Aβ42 fibrils^2, 5, 30^. It consistently showed that residue segments 17-20, 31-32, 34-36 and 39-41contributed mostly to the stability of Aβ42 fibrils. These segments overlapped mostly with β-strand-prone motifs gained from current MD simulations on dimers. Figure 6 also showed residue V12, V24 and residue 27-28 contributed favorably to a limited number of fibrils. Residue 27-28 overlaps with our MD observation as G25-Q27 is one of the β-strand-prone motifs. Figure 7 plotted two U-shaped Aβ42 fibril structures and two S-shaped Aβ42 fibril structures, with the hydrophobic β-strand-prone motifs from the current simulation highlighted in color. Significant diversities exist in the tertiary contacts of these motifs among U-shaped and S-shaped fibrils. For instance, L17-F20 (blue) locates closely with M35-V36 (yellow) and V39-V40 (red) in 2BEG, but in proximity with I31-I32 (purple) and M35-V36 (yellow) in 6TI6 and has contact with I31-I32 (purple), G35-N27 (black) in 2MXU and 5KK3. S-shaped fibrils usually display contacts between neighboring β-strand-prone segments at the C-terminal domain, probably determined by its fasting turning backbone nature at the loop region connecting two strands. This observation is consistent with the interstrand contacts observed in dimer 2 (figure 5). Therefore, the distinct dimers with short segments of β-strands developed in the current MD studies sampled the structural motif and similar tertiary contacts in these mature fibrils.

**Figure 6.**
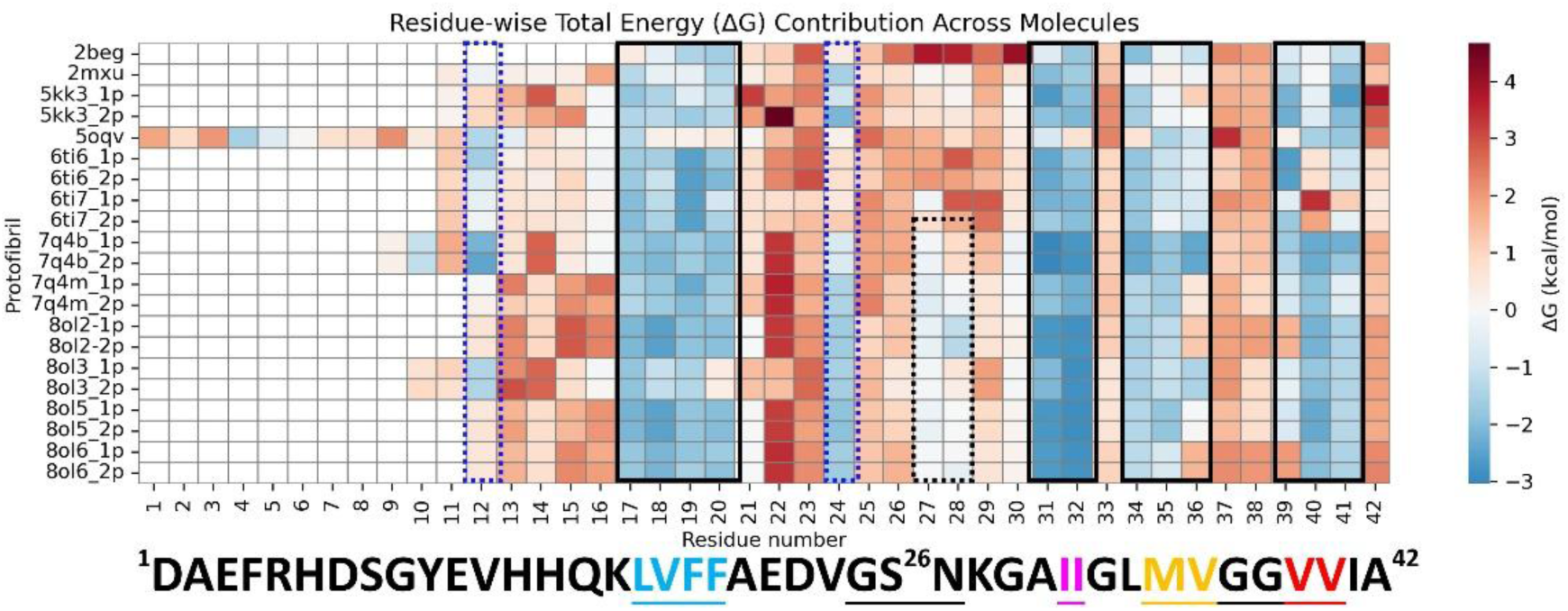
Individual residue contributions (residue numbers shown on the x-axis) to the stability of Aβ42 fibrils. If a fibril contains more than one protofilament, each protofilament is shown separately. The energy contribution was calculated using FoldX. The residues contributed mostly to stability are circled in black box with negative ΔG values, while the residues contributed moderately to stability or only favorably for some fibrils are circled in dotted rectangles.

**Figure 7.**
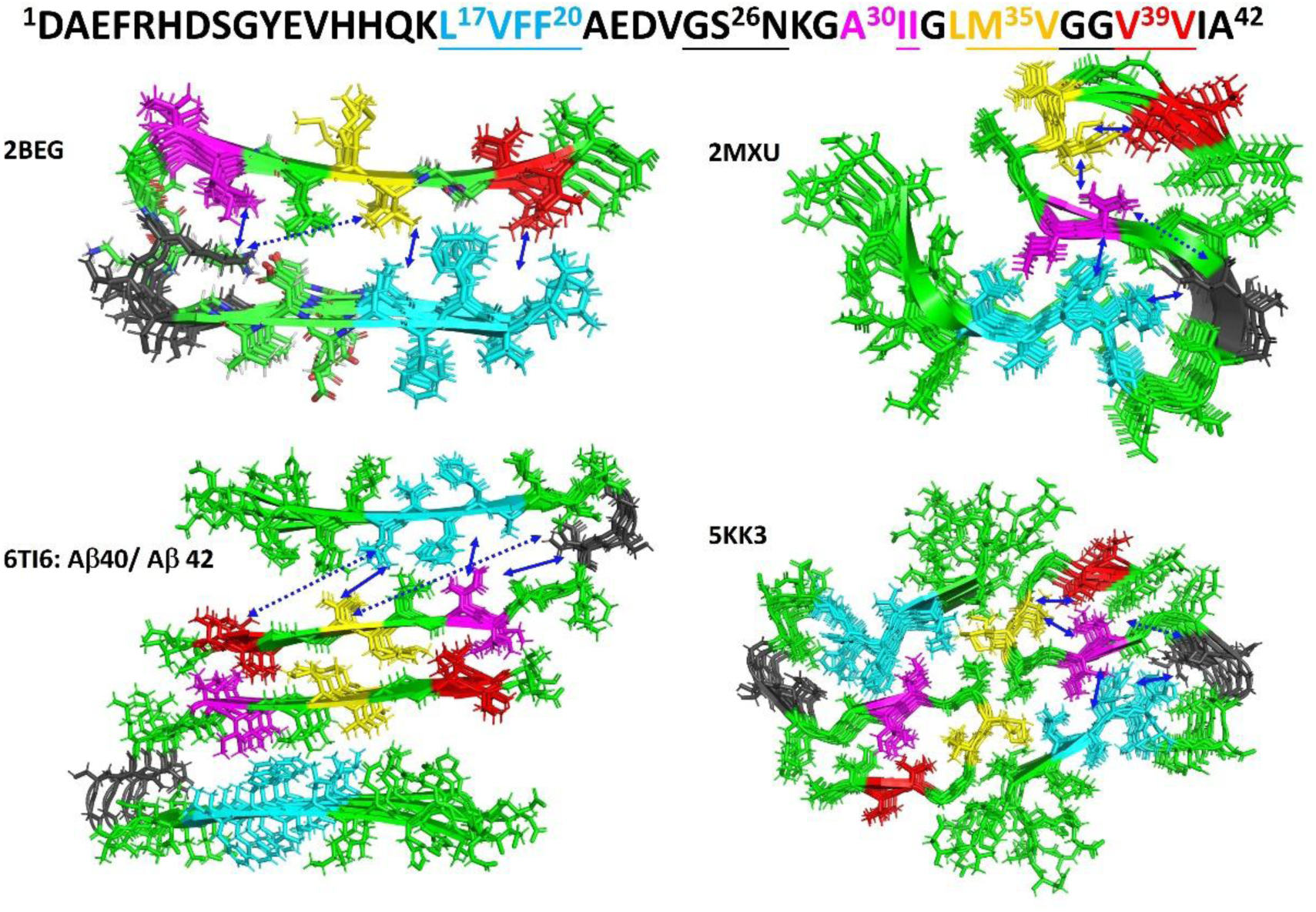
Representative Aβ42 fibril structures published, including U-shaped and S-shaped subunit structures. 6TI6 is formed by Aβ40 as well as Aβ42 sequence. All the other structures are formed by Aβ42. The sequence at the top line highlighted four hydrophobic segments and 1 polar segment in light blue, black, purple, yellow and red, corresponding to the colored stretches displayed in the PDB structures shown below. Figure is made by Pymol. The solid-line arrows indicate the space contacts while dotted-line arrows suggest no contacts with longer distances.

### 3.4 Effects of membrane bilayers on the nucleation of Aβ

A second insight from the current MD simulation is that the presence of membrane bilayer significantly affects the conformations of early-stage Aβ_42_ dimers. Our previous ssNMR study, which provided the inter-strand distances for the current MD simulation, investigated the structural changes in early-stage Aβ_42_ intermediates that bound to POPC bilayer. Since the experimental data was obtained on the membrane-bound aggregates (while small oligomers remain in solution were excluded in ssNMR sample preparation), represented by dimer 1, we compare its time-dependent evolution with the experimental trend. First, structural convergence for the N-terminal residues agreed with the closer contacts within residue 1-10 (Figure 5). We determined experimentally that different segments in Aβ42 sequence formed closer inter-strand distances at different rates, and the N-terminal residues converged at later stages compared to other segments. Noting that the ssNMR-based short inter-strand distances (e.g., 5.4 – 7 Å) were not applied to residues A2, F4 or G9, their proximities in dimer 1 developed at later stage of simulation may be driven by the cooperative fashion of structural convergence as shown in the ssNMR work. Second, the tertiary contacts between F19 and I32/L34 (Figure 5 and S4) were displayed at later stage of simulation, agreed with our two-dimensional (2D) ssNMR results with sensitive-enhanced dynamic nuclear polarization (DNP)^18^. Experimentally, these contacts were seen after six-hour incubation, accompanied by significant enhancement of the structural ordering of the membrane-bound Aβ_42_ intermediates, in prior to the onset of fibril elongation. This consistency implies that certain key tertiary interactions are crucial to produce stable and effective membrane-associated Aβ_42_ nuclei.

Additionally, the present MD simulation suggested that the formation of U-shaped conformation may help the early-stage Aβ42 dimers maintain in bilayers, while the dimers with extended conformation disassociated from membranes more easily. In the meanwhile, the membrane association was likely to reduce the secondary structural diversities in early-stage dimers compared to the one remaining in solution. We showed previously that the Aβ peptides added externally to membranes followed different pathways, depending on their membrane association. Membrane-bound Aβ nucleated more rapidly than those remaining in solution. And the membrane-associated mature Aβ fibrils from bilayers with different compositions showed uniform molecular structural features^31^. One explanation, based on the present simulation, is that the membrane association promotes the formation of structurally ordered nucleus by reducing the conformational flexibility in early-stage small oligomers.

## 4. Conclusions

In conclusion, our current MD simulations on the structural evolution of Aβ_42_ dimers demonstrate an approach to study early-stage disordered Aβ oligomers. Utilizing the ssNMR-based restraints, the structural properties and time evolutions of Aβ oligomers can be gleaned from the simulation more efficiently. The ssNMR-steered MD simulation could save computation power, holding potential for future investigations of Aβ aggregation with longer timescales. Within the current ~100-ns-simulation run, individual Aβ42 dimers showed distinct structural features and evolution pathways, depending on their association with membrane bilayers. Certain β-strand-prone motifs (i.e., hotspots) involving both hydrophobic and polar residues were identified, matching the structural features observed in the mature fibrils. The convergence and formation of higher order interactions between these motifs were influenced by environmental factors such as the close association of bilayers. Further MD simulation with the appropriate incorporation of ssNMR-based Aβ-lipid contact restraints with force are expected to shed light on the underlying structural basis.

## Author Contributions

A.L.C and B. S.L.C carried out the experiments, analyzed the data and prepared the figures. W.Q. wrote the main manuscript. W.Q. also conceived and supervised the project.

## Acknowledgements

We thank the Science and Engineering Fair organized by Thomas Jefferson High School for Science and Technology for encouraging our participation in research. We also thank Dr. Katherine Phillips and Ms. Emily Owens for their assistance. Parts of the coding for analyzing data were generated with the assistance of OpenAI ChatGPT (https://openai.com/index/chatgpt/).

