## supplemental files for "Molecular Dynamic simulations of Aβ42 dimers with solid-state NMR restraints capture the key structural motifs in Aβ42 fibrillation pathways"

**Supplementary Figures**

Figure S1. The secondary structure analysis using STRIDE algorithm in VMD. Light gray represents random coil (C), blue represents β-strand (E), green represents alpha helix (H), purple represents $3_{10}$-helix (G), orange represents π-helix (I), brown represents isolated bridge (B), pink represents turn (T) and dark gray represents bend (S). Dimer 1 formed by protein A and B developed β-strands at around residue 16-20. Dimer 2 formed by protein C and D developed β-strands at around residue 15-18 and residue 30-31. Dimer 3 formed by protein E and F did not display β-strands for a continuous period of time.

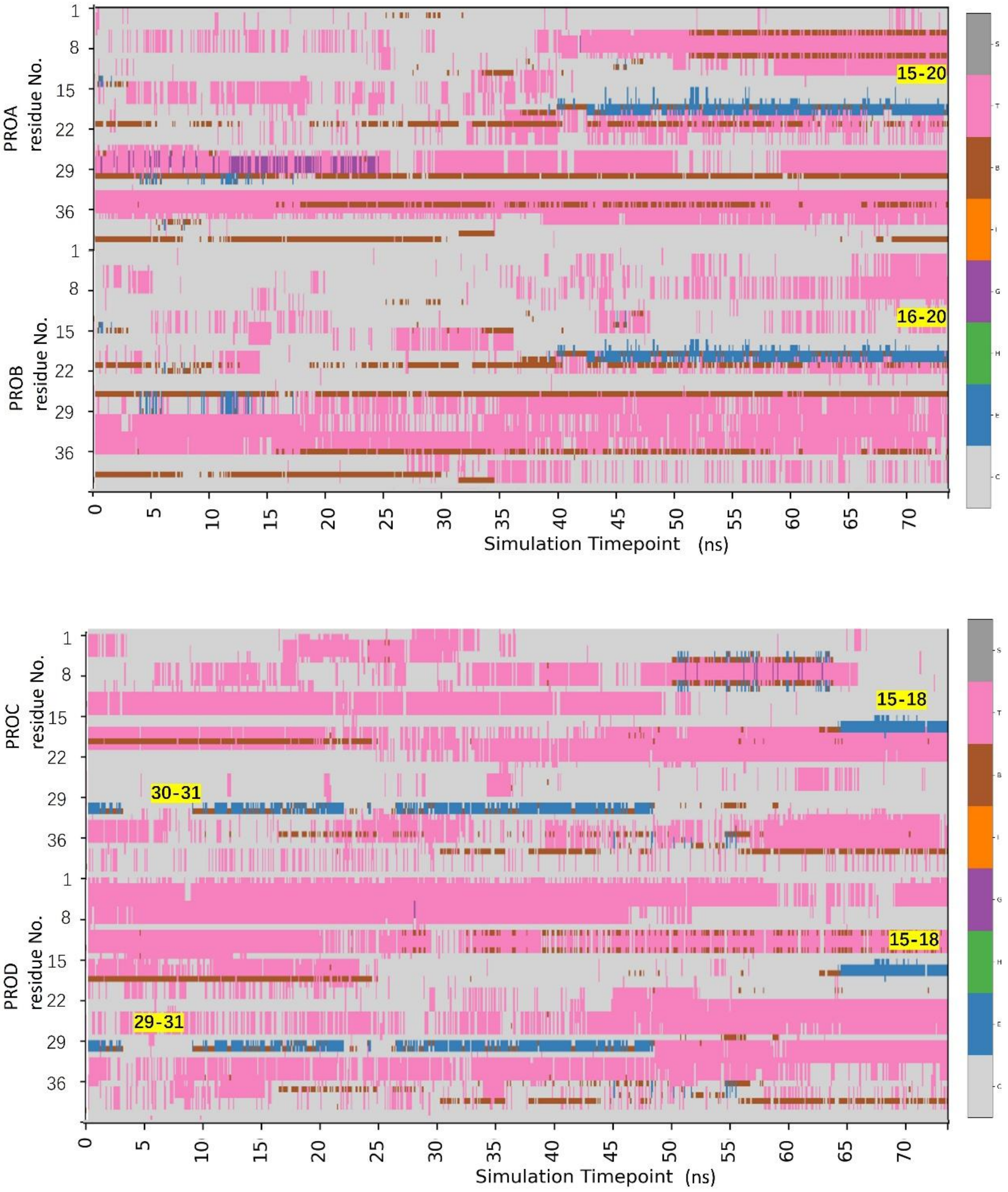

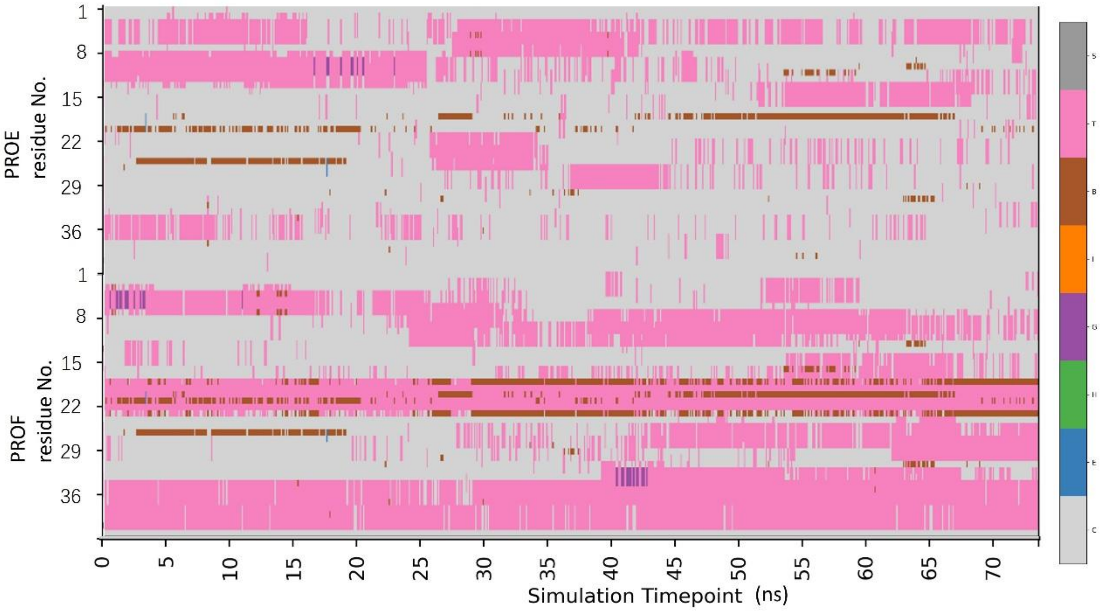
PROE residue No.
PROF residue No.
1
8
15
22
29
36
Simulation Timepoint (ns)
0
5
10
15
20
25
30
35
40
45
50
55
60
65
70

Figure S2. The secondary structure analysis of dimer 1 using STRIDE algorithm in VMD. Light gray represents random coil (C), blue represents β-strand (E), green represents alpha helix (H), purple represents $3_{10}$-helix (G), orange represents π-helix (I), brown represents isolated bridge (B), pink represents turn (T) and dark gray represents bend (S). Dimer 1 formed by protein A and B maintained β-strands at around residue 16-19 in POPC bilayers.

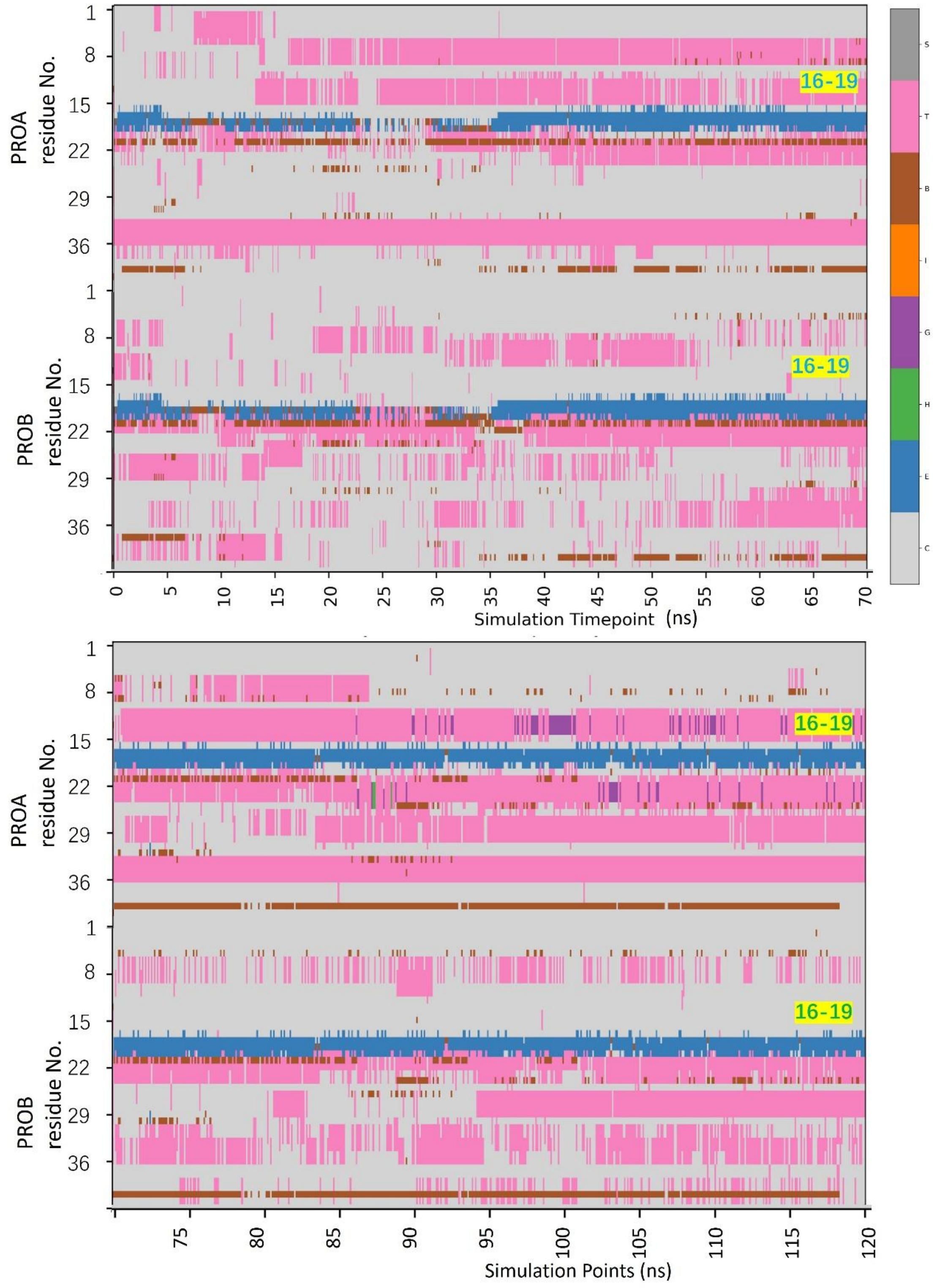

Figure S3. The secondary structure analysis of dimer 2 using STRIDE algorithm in VMD. Light gray represents random coil (C), blue represents β-strand (E), green represents alpha helix (H), purple represents $3_{10}$-helix (G), orange represents π-helix (I), brown represents isolated bridge (B), pink represents turn (T) and dark gray represents bend (S). Dimer 2 formed by protein C and D developed β-strands at many stretches in POPC bilayers, including at around residue 16-19.

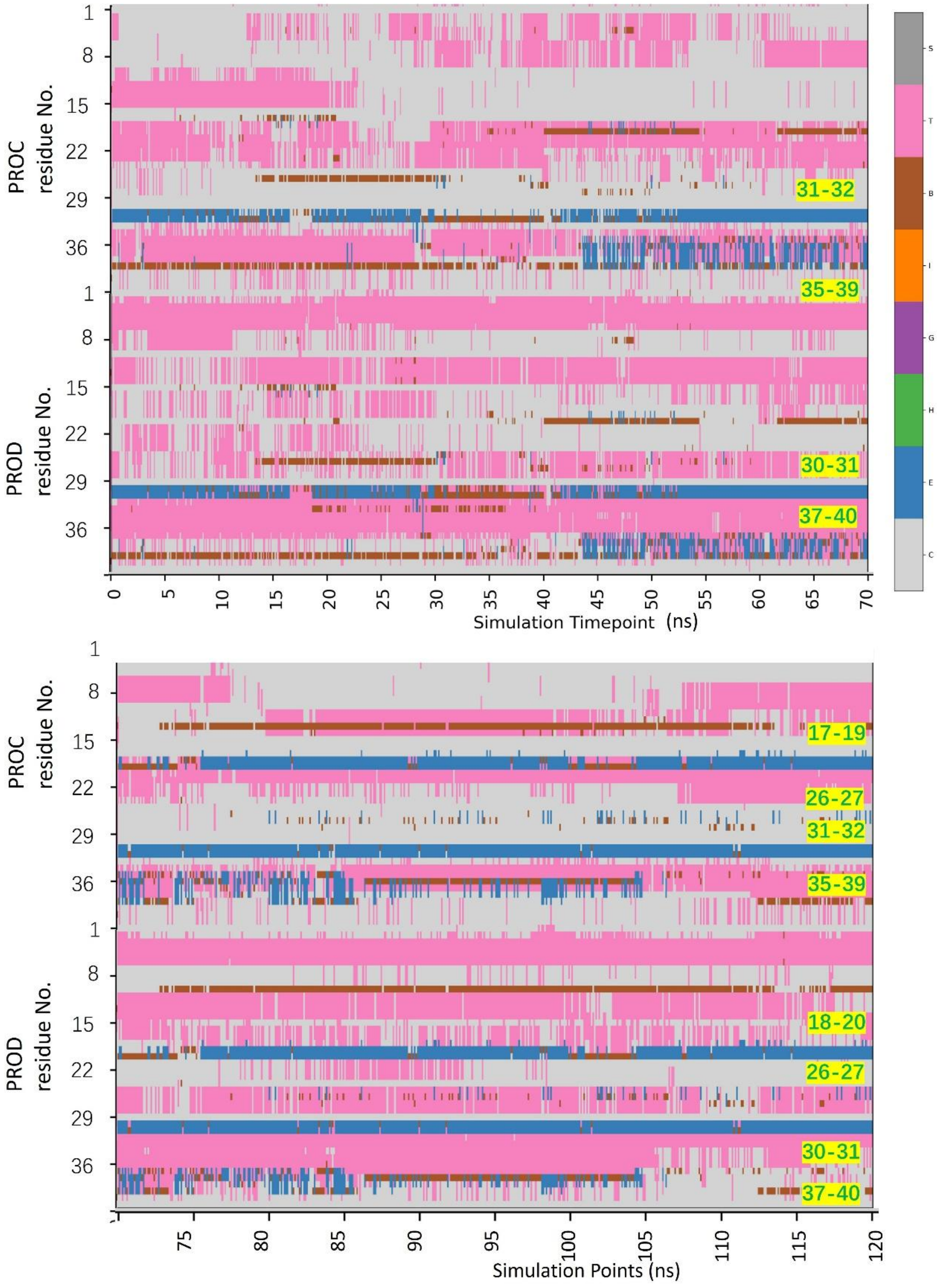

Figure S4. The distance variations between F19 and I32, and between F19 and L34 for dimer 1, the U-shape dimer, in the presence of POPC. The distance is calculated using the center of mass for each residue. Only the minimum distance is obtained. The distances are gradually decreased and stabilized in the period of 120ns MD simulation.

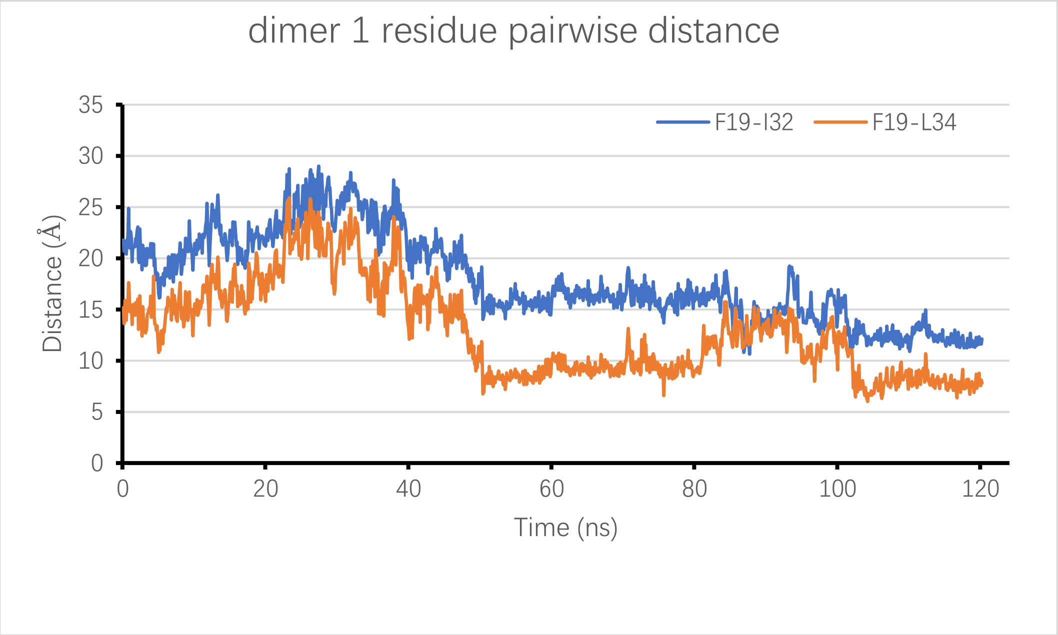
